# Miniature bioreactor arrays for modeling functional and structural dysbiosis in inflammatory bowel disease

**DOI:** 10.1101/2025.05.09.653173

**Authors:** Kira L. Newman, Alexandra K. Standke, Gabrielle James, Kimberly C. Vendrov, Naohiro Inohara, Ingrid L. Bergin, Peter D.R. Higgins, Krishna Rao, Vincent B. Young, Nobuhiko Kamada

## Abstract

Alterations in the gut microbiota, known as gut dysbiosis, are associated with inflammatory bowel disease (IBD). There is a need for model systems that can recapitulate the IBD gut microbiome to better understand the mechanistic impact of differences in microbiota composition and its functional consequences in a controlled laboratory setting. To this end, we introduced fecal samples from patients with Crohn’s disease (CD) and ulcerative colitis (UC), as well as from healthy control subjects, to miniature bioreactor arrays (MBRAs) and analyzed the microbial communities over time. We then performed two functional assessments. First, we evaluated the colitogenic potential of the CD microbiotas in genetically susceptible germ-free IL-10-deficient mice and found that colitogenic capacity was preserved in a bioreactor-cultivated CD microbiota. Second, we tested impaired colonization resistance against *Clostridioides difficile* in UC microbiotas using the MBRA system and found that UC microbiotas were innately susceptible to *C. difficile* colonization while healthy microbiotas were resistant, consistent with what is seen clinically. Overall, our results demonstrate that IBD microbiotas perform comparably to healthy donor microbiotas in the MBRA system, successfully recapitulating microbial structure while preserving IBD-specific functional characteristics. These findings establish a foundation for further mechanistic research into the IBD microbiota using MBRAs.

## Introduction

Alterations in the gut microbiota, known as gut dysbiosis, are associated with inflammatory bowel disease (IBD). Individuals with IBD exhibit reduced stability of the microbiota over time, decreased connectedness of the microbial community, and lower α-diversity compared to healthy controls. These changes may contribute to the pathogenesis of IBD and its complications.^1^ Gut microbiota features are predictive of the development of IBD in relatives of people with IBD. They also can predict recurrent *Clostridioides difficile* (*C. difficile*) infection in individuals with IBD and are associated with other complications, like fistulae and strictures.

There is a need for model systems that can recapitulate the IBD gut microbiome to better understand the mechanistic impact of differences in microbiota composition and its functional consequences in a controlled laboratory setting. One type of *in vitro* system that has been used to study *C. difficile* colonization in healthy microbiotas is bioreactors. These include single- and multi-chamber continuous flow systems that have helped elucidate the effects of strain competition in *C. difficile* infection and the impact of different antibiotic treatments on *C. difficile* persistence.^2-9^ A more recent development has been the use of higher-throughput simplified bioreactors capable of running multiple samples in parallel, also known as miniature bioreactor arrays (MBRAs).^10^ MBRAs have been shown to generate stable, reproducible microbiotas from healthy human fecal samples.^11^ They have been used to generate simplified microbial communities that resist *C. difficile* colonization, investigate the importance of amino acids for *C. difficile* colonization, test the effect of polyphenols on *C. difficile* susceptibility of the gut microbiome, and evaluate the effect of dietary emulsifiers on the healthy gut microbiome.^12-15^

Because of the altered diversity and composition of the gut microbiota in IBD, it remains unclear if the MBRA model, originally designed for cultivating a healthy gut microbiota, can reliably sustain stable and reproducible IBD patient-derived microbiotas while preserving their structural and functional characteristics. Prior applications of bioreactor systems to the cultivation of the IBD microbiota have used more complex three-stage and twin-vessel setups,^16, 17^ which offer lower throughput compared to MBRAs. Therefore, in this study, we sought to evaluate the robustness of the MBRA model for cultivating IBD microbiotas from both a taxonomic and functional standpoint. To this end, we cultured microbiotas from patients with Crohn’s disease (CD) and ulcerative colitis (UC), as well as from healthy control subjects, in MBRAs and analyzed the microbial communities over time. We then performed two functional assessments. First, we evaluated the colitogenic potential of the CD microbiotas in genetically susceptible germ-free IL-10-deficient mice. Second, we tested impaired colonization resistance against *C. difficile* in UC microbiotas using the MBRA system. Overall, our results demonstrate that IBD microbiotas perform comparably to healthy donor microbiotas in the MBRA system, successfully recapitulating microbial structure while preserving IBD-specific functional characteristics. These findings establish a foundation for further mechanistic research into the IBD microbiota using MBRAs.

## Results

### IBD-derived gut microbiotas can be stably and reproducibly cultivated in MBRAs

Following a similar protocol to Achtung, *et al*., we inoculated triplicate reactors with fecal samples from two donors with Crohn’s disease (CD) and two healthy control (HC) donors (**Figure 1A and 1B**). Samples were collected after 24 hours of acclimatization (time 0), at which point peristaltic flow was started with an 8-hour retention time. Samples were collected up to 9 days post-inoculation (**Figure 1B**). We assessed for microbial compositional changes using sequence analysis of PCR amplicons of the V4 region of the 16S rRNA gene sequencing from the initial inocula (donor stool samples) and samples from the bioreactors. We found that MBRAs supported diverse bacterial community growth from both HC and CD donor samples, as measured by the Shannon diversity index, inverse Simpson index, and number of species (i.e. richness) (**Figure 1C-E**). Shannon diversity and richness were stable from days 2-9 of the experiment (**Figure 1C-E**). Similar reduction in the overall number of species was seen in both HC and CD samples compared with the donor inocula, with an approximately 30-50% reduction (**Figure 1E**), similar to slightly lower than what has been previously reported for healthy donor samples cultivated in MBRAs^11^. The stability of richness and evenness were comparable between HC and CD donor samples by day 2 of cultivation (**Figure 1E and 1F**).

**Figure 1.**
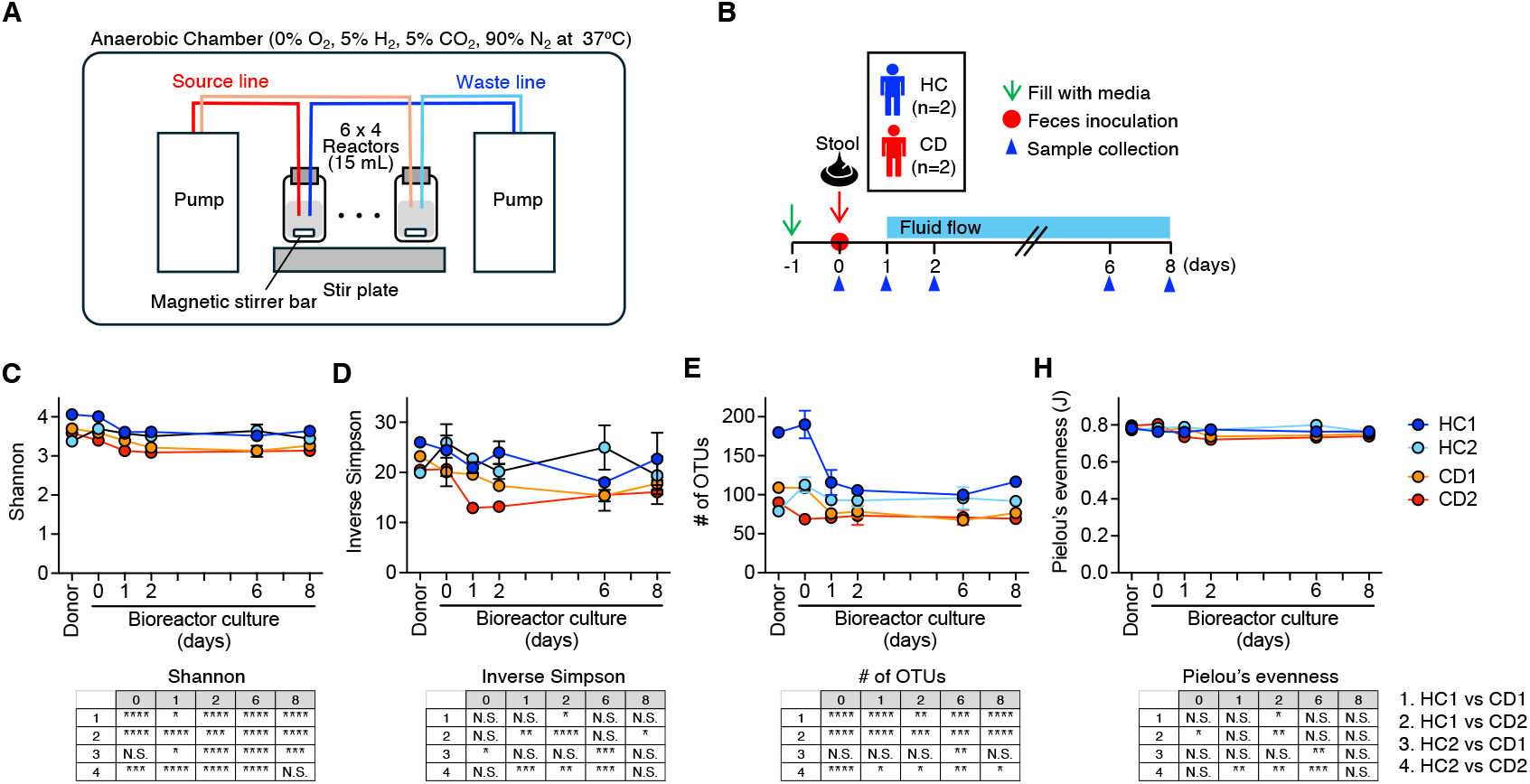
Miniature bioreactor arrays (MBRAs) allow for stable cultivation of healthy and inflammatory bowel disease (IBD) patient microbiomes. (A) MBRA setup inside an anaerobic chamber including six bioreactors per strip, each with an independent source and waste line attached to peristaltic pumps, stir plate for miniature stir bars, and sample port. Inset with magnified image of a single bioreactor. (B) Experimental schematic for testing stability of healthy and IBD microbiomes over time. (C-F) Measures of alpha-diversity in bioreactors over time based on 16S rRNA sequence data. Overall, Kruskal-Wallis p-values for differences between donors. Asterisks indicate time points with significant difference (p<0.05) between donors. All donors (n=2 healthy and n=2 IBD) with three technical replicates each.

### MBRA cultivation impacts community composition, but IBD microbiome remains distinct

We next evaluated the impact of cultivation in the MBRA on the structure of the microbial communities. We determined the relationships using Jaccard index and Bray-Curtis dissimilarity index. We visualized the relationships using non-metric multi-dimensional scaling (NMDS). As expected, each fecal community was distinct. Cultivation led to shifts in composition as measured by β-diversity indices compared to the donor inocula, but the communities remained separate from those derived from different donors (**Figure 2A and 2B**). There was a similarity between replicates from the same donor (**Figure 2A and 2B**). Adjustment to the MBRA environment led to significant decreases in the relative abundance of multiple families, including Bifidobacteriaceae, Lachnospiraceae, Lactobacillaceae, Ruminococcaeae, and Pasturellaceae (**Figure 2C**). There were also significant increases in the relative abundance of Bacteriodaceae, Peptostreptococcacus-Tissierellales, and Enterobacteriaceae (**Figure 2C**). These reflect some of the expected changes to a strictly anaerobic environment with limited carbohydrate availability.

**Figure 2.**
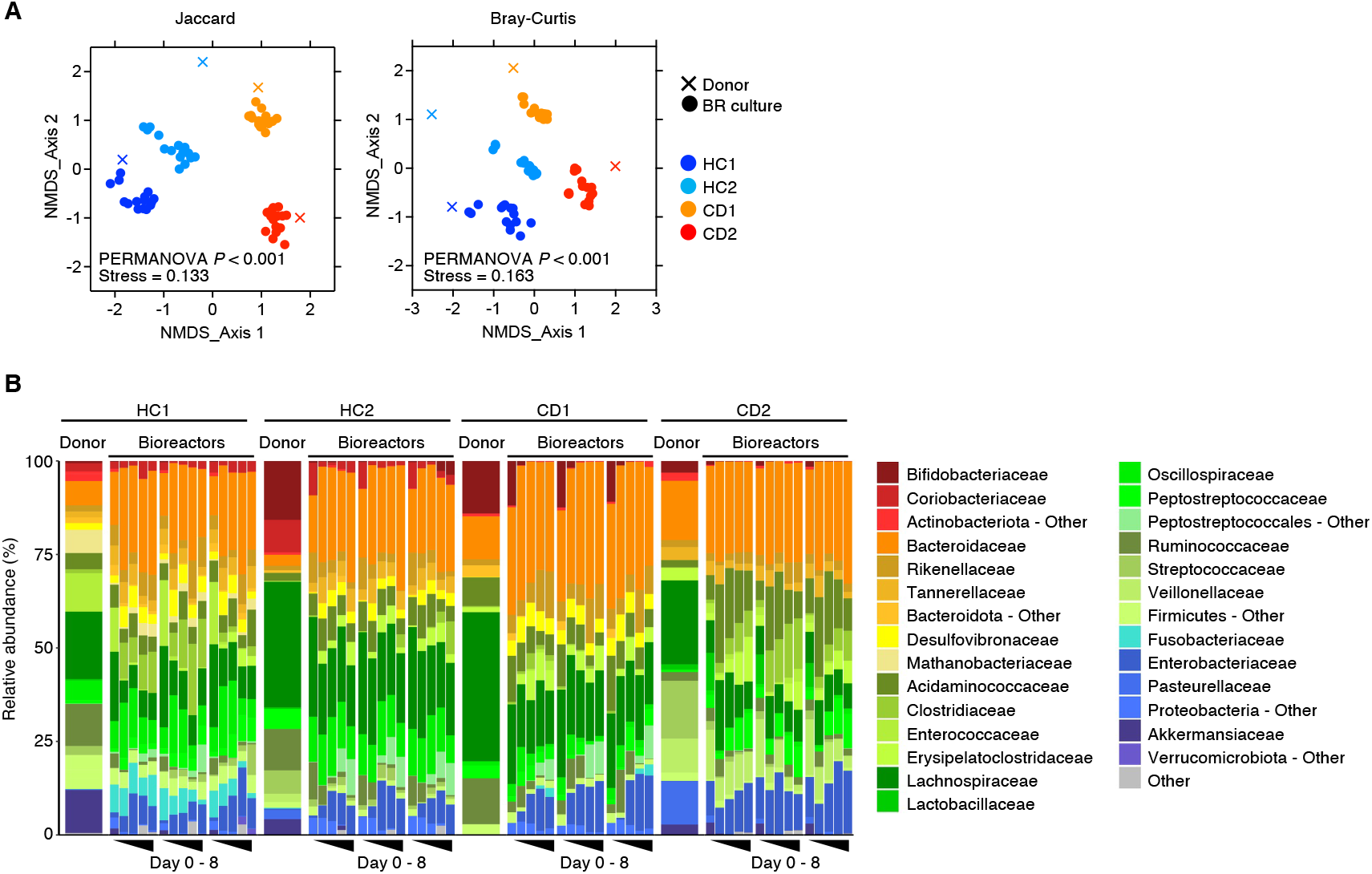
Beta-diversity in miniature bioreactor arrays (MBRAs) colonized with healthy and inflammatory bowel disease patient microbiomes is stable and donor-specific. (A and B) Two-dimensional NMDS plots summarizing beta-diversity measures of MBRA communities demonstrating clustering of points by donor (color coded). Triangles represent donor microbiome. Round dots represent samples from MBRAs (n=3 per donor) and time points (n=5). PERMANOVA comparing different donor groups. (C) Bacterial family-level composition. Bar plots show relative abundance (%) of bacterial families with at least 5% relative abundance in one or more samples. Samples are clustered by donor. Within each donor cluster, technical replicates (bioreactors) are clustered and depict relative abundance at each time point. The “Other” category represents bacteria from families in phyla with less than 5% relative abundance in any sample.

### IBD microbiota propagated in MBRAs retains colitogenic capacity

Human microbiota-associated (HMA) gnotobiotic animals have been widely used to study the functional role of human-derived microbiotas *in vivo*.^18-22^ We and others have previously reported that dysbiotic gut microbiota in IBD exhibit latent inflammatory potential. While colonization with IBD microbiota does not immediately induce colitis in healthy germ-free mice, it leads to severe colitis in hosts with genetic susceptibility (i.e., IL-10 deficiency) or when exposed to inflammatory insults, such as adoptive T cell transfer.^22-24^ Therefore, we tested whether IBD microbiotas propagated in MBRAs retained the same functional features as the original samples. First, we compared the microbiota structures in HMA mice and MBRAs (**Figure 3A**). To do this, donor microbiotas from one HC and two CD subjects were inoculated into germ-free (GF) wild-type C57BL/6 mice and allowed to reconstitute for 3 weeks (**Figure 3A**). The same donor microbiota samples were also propagated in MBRAs (**Figure 3A**). Notably, α-diversity, measured by the Shannon and Inverse Simpson indices, was comparable between microbiotas propagated in GF mice and MBRAs (**Figure 3B**). Moreover, β-diversity analysis showed that microbiotas from the same human donors propagated in GF mice and MBRAs were highly similar and retained significant differences between donors (**Figure 3C**). These results suggest that HC and CD microbiotas reconstituted in GF mice (HMA mice) and those propagated *in vitro* using MBRAs maintained structural similarity. Next, these *in vitro* and *in vivo* propagated human microbiotas were used for the colonization in GF *Il10*^-/-^ mice. Consistent with our previous report, HC microbiotas did not induce the development of colitis when colonized in GF *Il10*^-/-^ mice (**Figure 3D**). In contrast, we previously reported that certain CD microbiotas exhibited colitogenic potential when colonized in GF *Il10*^-/-^ mice.^23^ In this experiment, we tested microbiotas from two different CD donors. Notably, one of these donor microbiota displayed a robust colitogenic capacity in GF *Il10*^-/-^ mice, as indicated by elevated fecal lipocalin 2 (Lcn2) levels and histological inflammation (**Figure 3D-F**). Next, we examined whether *in vitro* propagated microbiotas using MBRAs maintain the colitogenic capacity. Similar to the microbiota reconstituted in HMA mice, MBRA-propagated HC and non-colitogenic CD microbiotas did not induce colitis in GF *Il10*^-/-^ mice (**Figure 3D-F**). In contrast, the colitogenic CD microbiota retained its colitogenic capacity after propagation in MBRAs (**Figure 3D-F**). Thus, *in vitro* propagation in MBRAs effectively preserves colitogenic pathobionts present in CD microbiotas.

**Figure 3.**
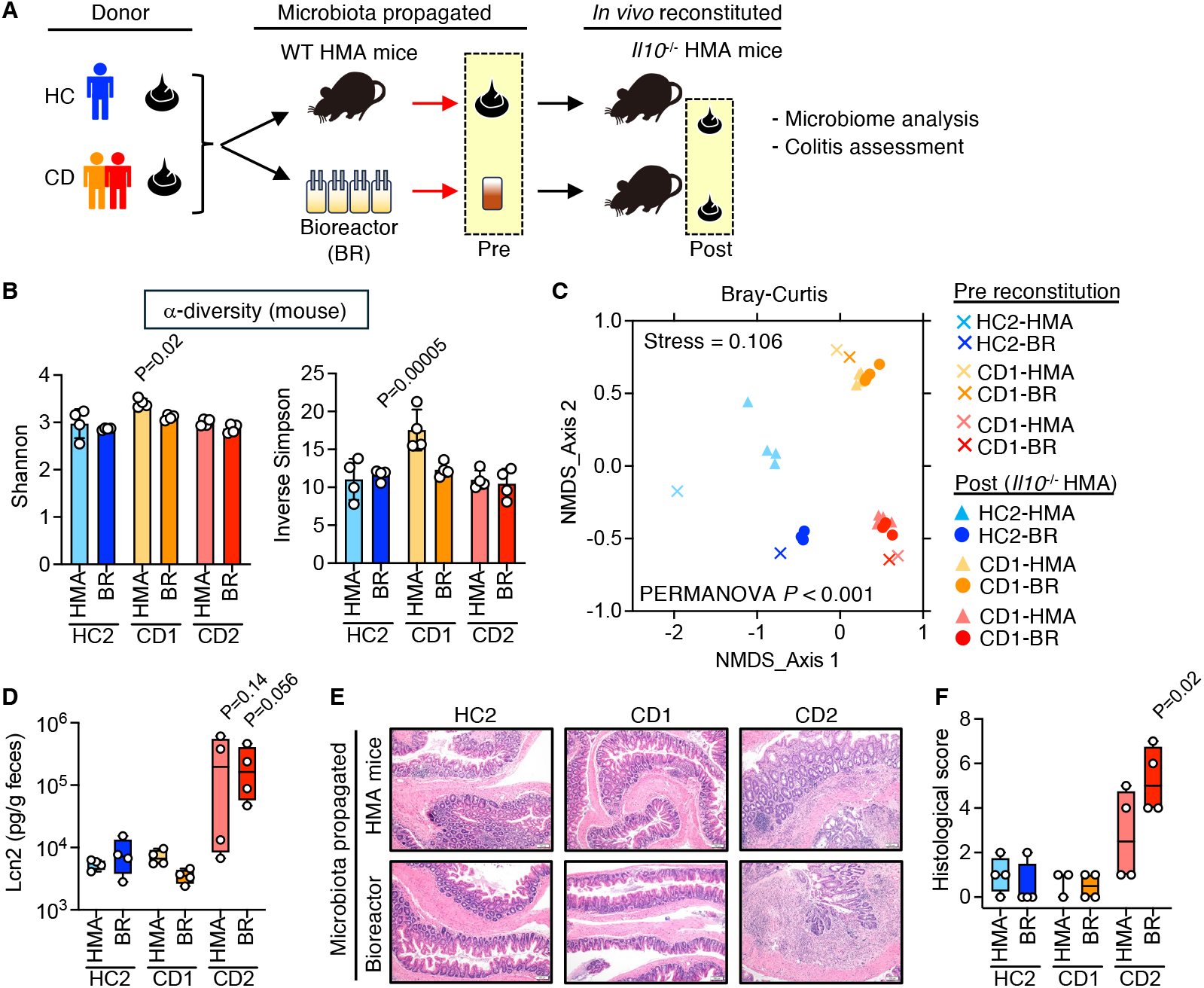
Bioreactors preserve colitogenic capacity of the Crohn’s disease microbiome in a human microbiota-associated (HMA) mouse model. (A) Experimental schematic illustrating comparison of standard IL10-knockout (KO) mouse HMA model and bioreactor-derived microbiome variation. For the standard model, human donor stool is inoculated into a wildtype germ-free mouse to generate an HMA mouse. This mouse’s colon and cecum contents are then collected after sacrifice and used to inoculate an IL10KO mouse. In the bioreactor-derived variation, human donor stool from the same donors is cultured in a bioreactor. The bioreactor-cultured microbiome is then used to inoculate an IL10KO mouse. Mice are then sacrificed and extent of colitis is assessed. (B) The alpha diversity of the microbiome based on 16S rRNA analysis of colon contents is similar between HMA and BR mice from the same donors. Diamonds indicate alpha-diversity measure for the human donor sample. Circles indicate individual mice. n=4 mice per group. (C) Two-dimensional NMDS plot of Bray-Curtis dissimilarity index. Circles indicate donor inocula (either standard HMA-derived or bioreactor-cultivated). Triangles indicate mice. Points are color coded by human donor. (D) Bar graph of fecal lipocalin 2 levels from mice in each group. (E) Representative cecal histology from mice in each group. (F) Histologic score of cecal inflammation. Each dot represents an individual mouse. Results are mean +/-standard deviation.

### MBRA-propagated IBD microbiota display impaired colonization resistance to an enteric pathogen

Next, we investigated another functional feature of IBD microbiota using MBRA-propagated stool samples, specifically examining susceptibility to the enteric pathogen *C. difficile*. It has been reported that IBD, particularly in UC patients, is associated with an increased risk of enteric pathogen infections, including *C. difficile*.^25^ Previously, we demonstrated that UC microbiota-colonized HMA mice exhibit impaired colonization resistance, allowing *C. difficile* to colonize and proliferate. Consequently, *C. difficile* infection in these mice leads to lethal colitis.^18^ To assess potential impairments in UC microbiota resistance to *C. difficile* colonization, fecal samples from HC and UC donors (n = 6 and 4, respectively) were propagated in MBRAs (**Figure 4A and Supplemental Figure 1A-D**). After a 3 day stabilization period of the microbiota in MBRAs, *C. difficile* was inoculated *in vitro* (**Figure 4A**). When assessed after stabilization, α-diversity, measured by the Shannon and Inverse Simpson indices, was lower in UC vs HC microbiota in the MBRAs, though not in the initial donor samples. However, β-diversity analysis showed that UC and HC microbiotas propagated in MBRAs were significantly different between donors (PERMANOVA comparing MBRA samples between donors p = 0.005) (**Figure 4B, C**). Consistent with findings in HMA mice, the majority of HC microbiotas were resistant to *C. difficile* colonization, with inoculated *C. difficile* showing little to no growth in HC microbiota within the MBRAs (**Figure 4D**). In contrast, inoculated *C. difficile* proliferated extensively in most IBD microbiotas (**Figure 4D**). Thus, *in vitro* propagated IBD microbiotas in MBRAs effectively replicate the impaired colonization resistance against *C. difficile* observed in IBD patients.

**Figure 4.**
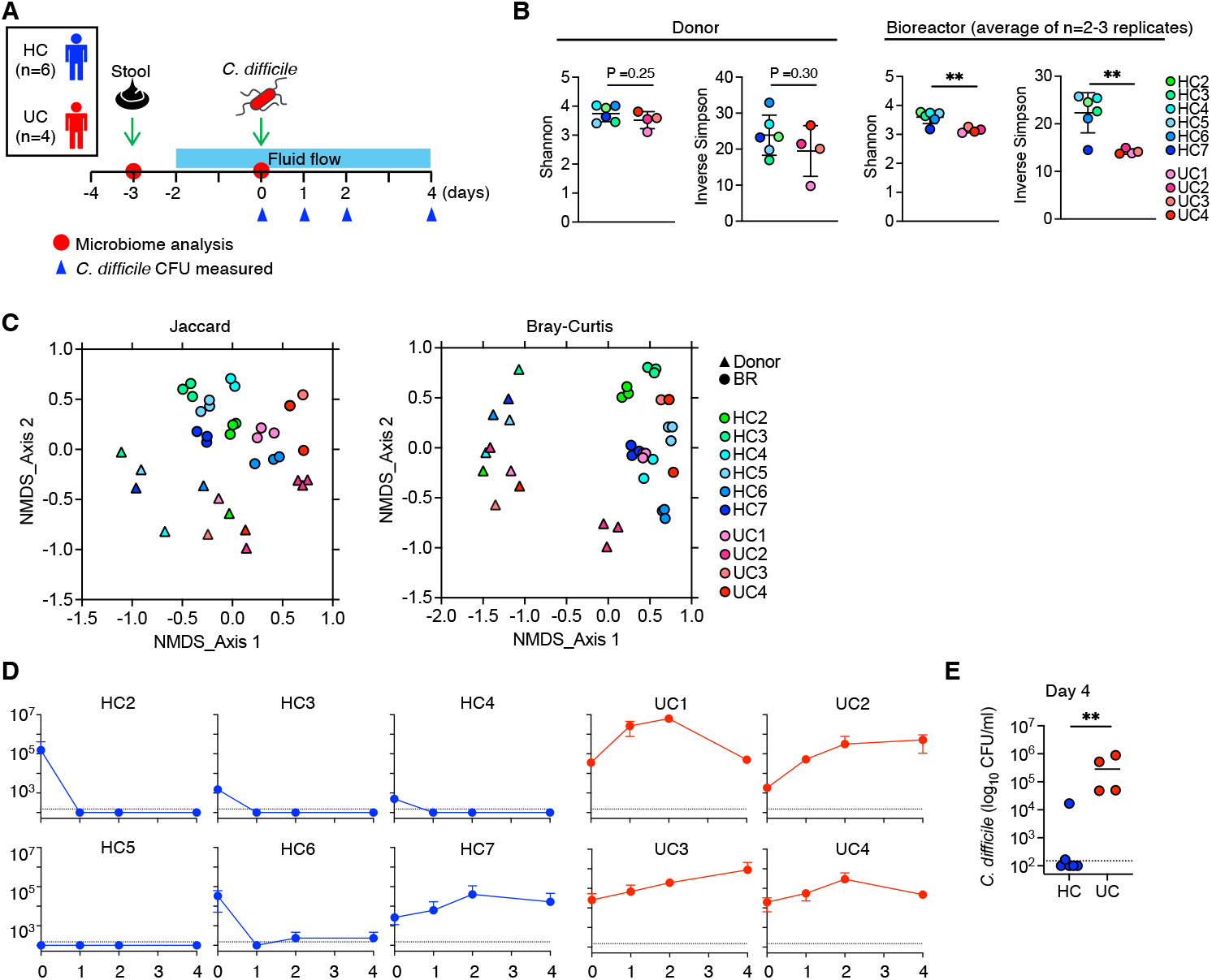
Bioreactors preserve susceptibility to C. difficile colonization of the ulcerative colitis (UC) microbiome. (A) Experimental schematic illustrating colonization of bioreactors in the miniature bioreactor array (MBRA) with human donor samples (n=6 healthy and n=4 UC). After one day to establish the community, peristaltic flow of media is begun on Day -2. After 48 hours, communities are then challenged with inoculation of vegetative cells of *C. difficile* strain VPI 10463 on Day 0. Samples are then collected over the subsequent four days to assess for colonization. (B) Alpha diversity of microbiome based on 16S rRNA analysis of donor and Day 0 MBRA samples. (C) Beta diversity illustrated with two-dimensional NMDS plots of Jaccard index and Bray-Curtis dissimilarity index. (D) Graphs of colony-forming units (CFU) of *C. difficile* from bioreactors at each time point. We performed three technical replicates per sample. Data are presented as mean +/-standard deviation. (E) Comparison of the CFU of *C. difficile* at day 4 post-inoculation. Each point represents the mean CFU for each donor. **; p <0.01 by Wilcoxon rank-sum test.

## Discussion

In this study, we demonstrated that IBD microbiotas can be effectively recapitulated using MBRAs - both structurally and functionally. MBRA-propagated IBD microbiotas exhibited a microbial composition comparable to that of reconstituted HMA mice. Beyond structural recapitulation, IBD microbiotas propagated in MBRAs retained their functional characteristics, including latent colitogenic capacity and impaired colonization resistance to *C. difficile*. Since these are well-established functional traits of IBD-associated dysbiotic microbiotas, the MBRA system represents a powerful tool for studying IBD microbiota function.

Due to various innovations in computational biology, omics technologies, including microbiome analysis, have advanced rapidly. In the field of IBD research, multi-omics approaches have advanced microbiome science from correlative studies toward more functional and mechanistic investigations. Several studies have reported that gut dysbiosis in IBD patients is not merely an imbalance in bacterial structure. Rather, the inferred functions of the microbiota are also perturbed in patients with IBD.^26-29^ However, while multi-omics approaches strongly indicate functional disturbances in the microbiota, biological models are still required to validate these proposed functional changes with greater precision and mechanistic insight. In this context, gnotobiotic animal models remain the gold standard for assessing the function of individual human-derived bacterial strains, bacterial consortia, or the entire microbiota.^30^ Indeed, we and others have employed the human microbiota-associated (HMA) gnotobiotic mouse model and successfully recapitulated key functional traits of dysbiotic microbiota in IBD patients, such as colitogenicity and susceptibility to *C. difficile*. ^18, 23^

However, the HMA gnotobiotic system is low-throughput and costly. Also, only a limited number of researchers with access to germ-free/gnotobiotic facilities can conduct this type of research. In this regard, the present study demonstrates that the MBRA system can recapitulate the altered structure of IBD-associated microbiota, comparable to HMA mice. Beyond structural replication, the MBRA-propagated microbiota also retains the functional defects observed in IBD patients. Thus, the MBRA system serves as a valuable adjunct or alternative to HMA mice for studying functional defects in IBD-associated microbiotas. Also, not only for cost reasons, there is a need for validated non-animal and animal-sparing models for microbiome research for ethical reasons, including the 3R European rules.^31^ Together, the MBRA represents a powerful open-source option with proven use in multiple labs across different continents.^14, 32, 33^

Furthermore, the MBRA system offers certain scientific advantages over the HMA mouse model. For example, it allows for the study of site-specific microbiota functions from different intestinal regions, independent of the effects of microbes from other sites (e.g., fecal versus mucosal).^34^ Moreover, the MBRA system enables the study of the direct impact of nutrients, drugs, chemicals, or other modifications on the microbiota while eliminating secondary effects from host-related changes induced by these interventions. This ability to focus on microbe-microbe interactions in the absence of host factors can allow for identification of microbiome functions for which host-specific inputs are not necessary.

Also, IBD presents with a diverse range of clinical phenotypes and a major challenge in IBD care is developing personalized strategies for treatment. For example, some UC patients experience reduced inflammation following fecal microbiota transplantation (FMT),^35, 36^ while others do not respond as effectively.^37^ These variations in FMT efficacy may be attributed to differences in the baseline composition of the gut microbiota or challenges in identifying optimal donor-recipient matches. Paired with an improved understanding of the specific features of a dysbiotic gut that associate with efficacy, further studies could use the MBRA system to screen for the most effective donor in correcting dysbiosis before its application in a clinical setting.

Taken together, our work supports the MBRA system as a valuable tool for studying the functions of dysbiotic microbiota in IBD, as we as for the future implementation of personalized medicine in IBD.

## Materials and Methods

### Fecal sample collection and preparation

Fecal samples were collected from 12 individuals who had not been treated with oral antibiotics within the previous three months. All participants provided consent to participate in the study.

Protocols for collection and use of fecal samples were reviewed and approved by the Institutional Review Board at the University of Michigan (HUM00041845). Detailed patient information is provided in **Table S1**.

Samples were self-collected by participants in commode specimen containers, sealed in a plastic bag, stored at 4 ºC, and returned to the lab within 24 hours. Fecal samples were aliquoted and frozen at -80 ºC until later resuspended in reduced phosphate buffered saline (PBS) under anaerobic conditions (Coy Anaerobic Chamber with 5% H_2_, 5% CO_2_, 90% N_2_ atmosphere). For samples used in bioreactor experiments, they were suspended at 25% w/v in PBS, per protocol by Achtung, *et al*..^10^ These were then vortexed at >2,500 rpm for 5 minutes, centrifuged at 200 g for 5 min, and the supernatants immediately used or preserved at -80 ºC for further analysis. For samples used in murine experiments, they were suspended in reduced PBS at a concentration of 1 mL PBS per 0.1 g feces. These samples were then strained through a 100 μm strainer basket (Fisher) and immediately used or preserved at -80 ºC for further analysis.

### Bioreactor experiments

As described previously, MBRAs were partially assembled, autoclaved, and then transferred to an anaerobic chamber (5% H_2_, 5%CO2, 90%N_2_). In the anaerobic chamber, the MBRAs were assembled and the chamber was warmed to 37 ºC. Each MBRA was configured to hold 15 mL volume. They each contain a magnetic stir bar, sample port, input port for slow continuous delivery of fresh media, and waste port for removal of excess volume. They are configured with six MBRAs per 3D-printed strip and placed over a stir plate (**Supplemental Figure 2**). All experiments used bioreactor medium (BRM), prepared per published protocol.^10^ This media is low in fermentable carbohydrates and uses protein as the primary nutritional source. It is intended to mimic the nutritional conditions of the distal colon and has been used in prior MBRA experiments. MBRAs were connected under sterile conditions, filled with 15 mL of anoxic BRM, left to sit for 24 hours to confirm sterility, and then inoculated with fecal suspensions at 1% w/v final concentration. These were allowed to grow for 24 hours with gentle stirring prior to the initiation of peristaltic flow of media at 1.875 mL/hr. For *C. difficile* challenge experiments, communities were allowed to equilibrate for 48 hours prior to inoculation with ∼ 1x 10^5^ CFU/mL of vegetative cells of *C. difficile* VPI 10463. CFUs were then enumerated over time. Samples were collected from bioreactors using a sterile needle and syringe. Aliquots of samples were used for *C. difficile* enumeration by serial dilution and plating on selective agar (CHROmagar). Samples were then frozen at ≤-20 ºC.

### Human microbiota-associated (HMA) gnotobiotic mice

All animal husbandry was performed by the University of Michigan Germ-Free Facility. Mice were housed at the University of Michigan Germ-Free Facility in positive-pressure, individually-ventilated cages (ISOcage P; Techniplast, West Chester, PA) to maintain germ-free (GF) status and prevent cross-contamination during experiments. All mice were fed a sterilized rodent breeder diet 5013 (LabDiet, St. Louis, MO). Male and female mice aged 9-12 weeks were used in all experiments. All animals were handled in accordance with the protocols reviewed and approved by the Institutional Animal Care and Use Committee. As previously described, GF wild-type C57BL/6 mice were gavaged with 200 μl each of human-derived fecal inoculum and establish human microbiota-associated (HMA) gnotobiotic mice.^23^ HMA mice were then sacrificed after 21 days of microbiota reconstitution, and their cecum contents were kept frozen at -80 ºC until use. To assess the colitogenic capacity of the microbiota, frozen aliquots of HMA mouse-derived cecal contents were thawed in an anaerobic chamber, suspended 1:10 in PBS, and strained using a 100 μm filter basket. This HMA cecal suspension was used to inoculate into GF *Il10*^-/-^ C57BL/6 mice (200 μl per mouse). Alternatively, frozen stocks of bioreactor-processed human microbiota were thawed, and undiluted suspensions were inoculated into GF *Il10*^-/-^ C57BL/6 mice (200 μl per mouse). Human microbiota-reconstituted GF *Il10*^-/-^ mice (*Il10*^-/-^ HMA mice) were kept for 21 days. At 21 days, *Il10*^-/-^ HMA mice were sacrificed for the evaluation of colitis. Cecum and colon tissues were collected and fixed with 4% paraformaldehyde. Fixed tissues were processed, embedded, sectioned, and stained with hematoxylin and eosin (H&E) by the University of Michigan Histology Core. Slides were deidentified and submitted for blinded histological scoring by a veterinary pathologist at the University of Michigan Unit for Laboratory Animal Medicine In-Vivo Animal Core. Colon contents were suspended 1:10 in PBS then frozen at -20 ºC prior to submission to the University of Michigan Immunology Core for ELISA assay to quantify fecal lipocalin-2 (Lcn2) levels.

### Microbial community analysis by 16S rRNA gene sequencing

DNA was extracted from samples using the Qiagen Blood and Tissue kit according to instructions with the following modifications. Either 200 μl of suspended inoculum, liquid bioreactor sample, or one fecal pellet were used for DNA extraction. Samples were disrupted using beadbeating for 1.5 minutes with PowerBeads (Qiagen) on a Mini-Beadbeater-16 (BioSpec Products, Inc, Bartlesville, OK). The amount of buffer ATL used in the initial steps of the protocol was increased from 180 to 400 μl. The volume of proteinase K was increased from 20 to 40 μl. The amount of buffer AE used to elute DNA at the end of the protocol was decreased from 200 to 75 μl. The V4 region of bacterial 16S rRNA was sequenced using the Illumina MiSeq at the University of Michigan Microbiome Core/Host Microbiome Initiative.

Fastqs were processed by Qiime2 (version 2024.5),^38^ denoising with DADA2,^39^ mapping sequences against SILVA (release 138.1)^40-42^ using the q2-feature-classifier,^43^ and clustering ASVs at 100%. Shannon diversity index, inverse Simpson dissimilarity, and ASV richness were used to show community diversity. To compare the diversity between communities (i.e. β-diversity), we used the Jaccard similarity index and Bray-Curtis Dissimilarity. Bray-Curtis calculations were shown as a non-metric multidimensional scaling (NMDS) plot using the R package vegan (version 2.6-10; https://cran.r-project.org/web/packages/vegan/vegan.pdf).

### *C. difficile* cultivation

Spore stock of *C. difficile* strain VPI 10463 (ATCC 43255, Gift of Vincent Young) was prepared as previously described.^44^ Vegetative *C. difficile* was cultured in an anaerobic chamber (Coy Laboratory products) by streaking spores onto a pre-reduced selective agar plate (CHROmagar) then incubating overnight at 37 ºC. The next day a colony was subcultured into reduced BRM. After overnight incubation at 37 ºC, the culture was diluted 1:10 in reduced BRM then incubated at 37 ºC. An aliquot was monitored for growth using a plate reader (Tecan) for OD_600_. When OD_600_ was between 0.08–0.15, the culture was diluted 1:20 in reduced BRM then used to inoculate the MBRAs as above.

### Data visualization and statistical analysis

Figures and statistical analysis were generated in R (v.4.4.0) or GraphPad Prism (v10.0). Statistical tests used for the analysis of data are identified in the legend of each figure. Differences of *P* < 0.05 were considered significant.

## Acknowledgments

The authors thank the University of Michigan Microbiome Core, the Germ-Free Mouse Facility, the Advanced Genomics Core, and the ULAM Pathology Core (RRID:SCR_018823). Yadong Mao for technical assistance. This work was supported by the National Institute of Health DK134043 (K.L.N.), DK108901 (N.K.), DK119219 (N.K.), AI162787 (V.B.Y), AI124255 (V.B.Y.), and by the American College of Gastroenterology (K.L.N.).

## Authorship Contributions

K.L.N. and N.K. conceived and designed experiments. K.L.N. and G.J. conducted most of the experiments with help from A.K.S. and K.V.. N.I. performed microbiome analyses. I.L.B. performed pathology analyses. P.D.R.H., K.R., and V.B.Y. helped with critical advice and discussion. K.L.N. and N.K. analyzed the data. K.L.N. and N.K. wrote the manuscript with contributions from all authors.

## Data Availability

The microbiome data in this study are available at the NCBI Sequence Read Archive under BioProject (will be available before publication).

