## Supplemental Figure 1 and 2, Supplemental Table 1 for "Miniature bioreactor arrays for modeling functional and structural dysbiosis in inflammatory bowel disease"

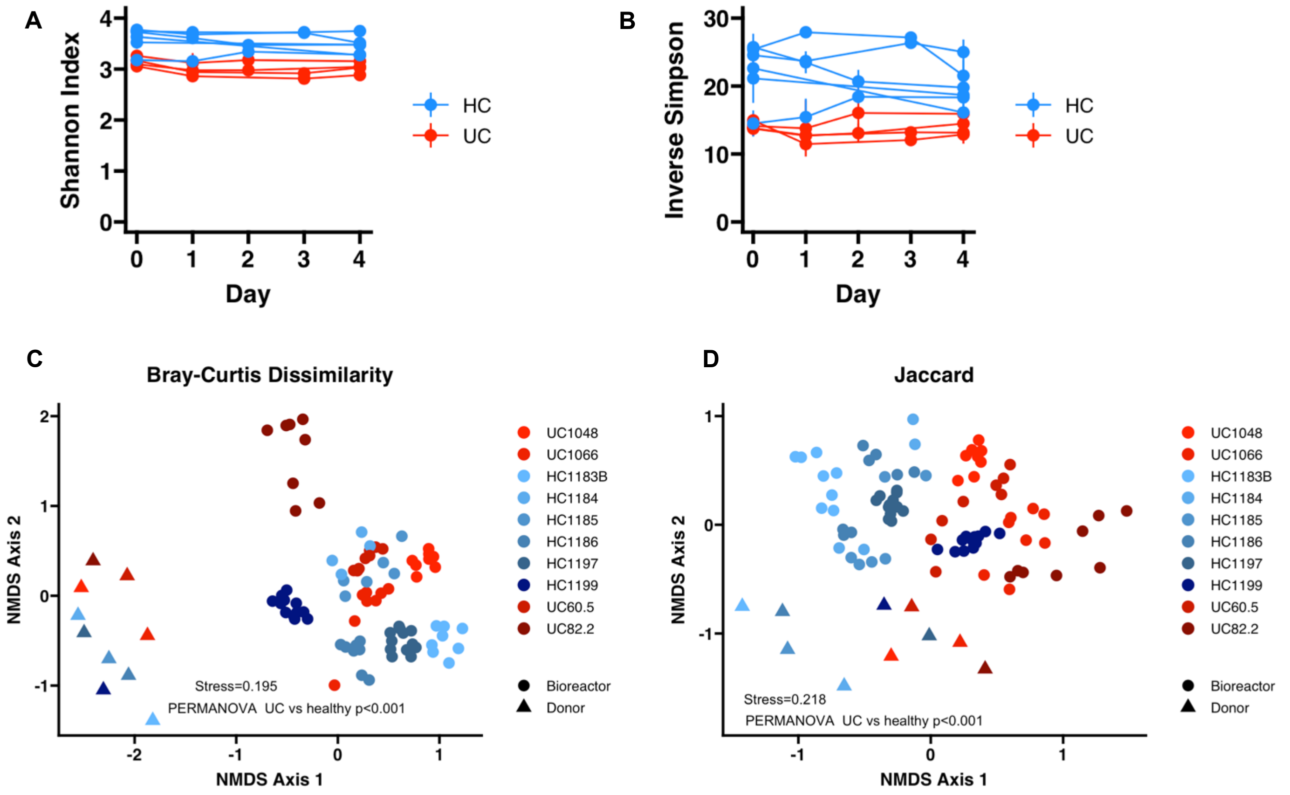


**Supplemental Figure 1.** Alpha and beta diversity of MBRA-cultivated HC and UC samples before *C. difficile* inoculation and after at all time points tested. (A-B) Alpha diversity of UC and HC samples grown in MBRAs remains stable before and after inoculation with *C. difficile* spores. Each line represents the mean +/- standard deviation alpha diversity for all bioreactors colonized with the same donor sample. n=1-3 replicates per time point per donor. No significant difference between Day 0 and Day 4 alpha diversity measures by Kruskal-Wallis tests. (C-D) Beta diversity of UC and HC samples grown in MBRAs remains stable after inoculation with *C. difficile* spores as illustrated by clustering by donor of all MBRA samples from all days tested. UC and healthy donor samples remain significantly different even after accounting for additional time points. n=1-3 replicates per time point per donor. PERMANOVA used for significance testing between all UC versus all HC samples.

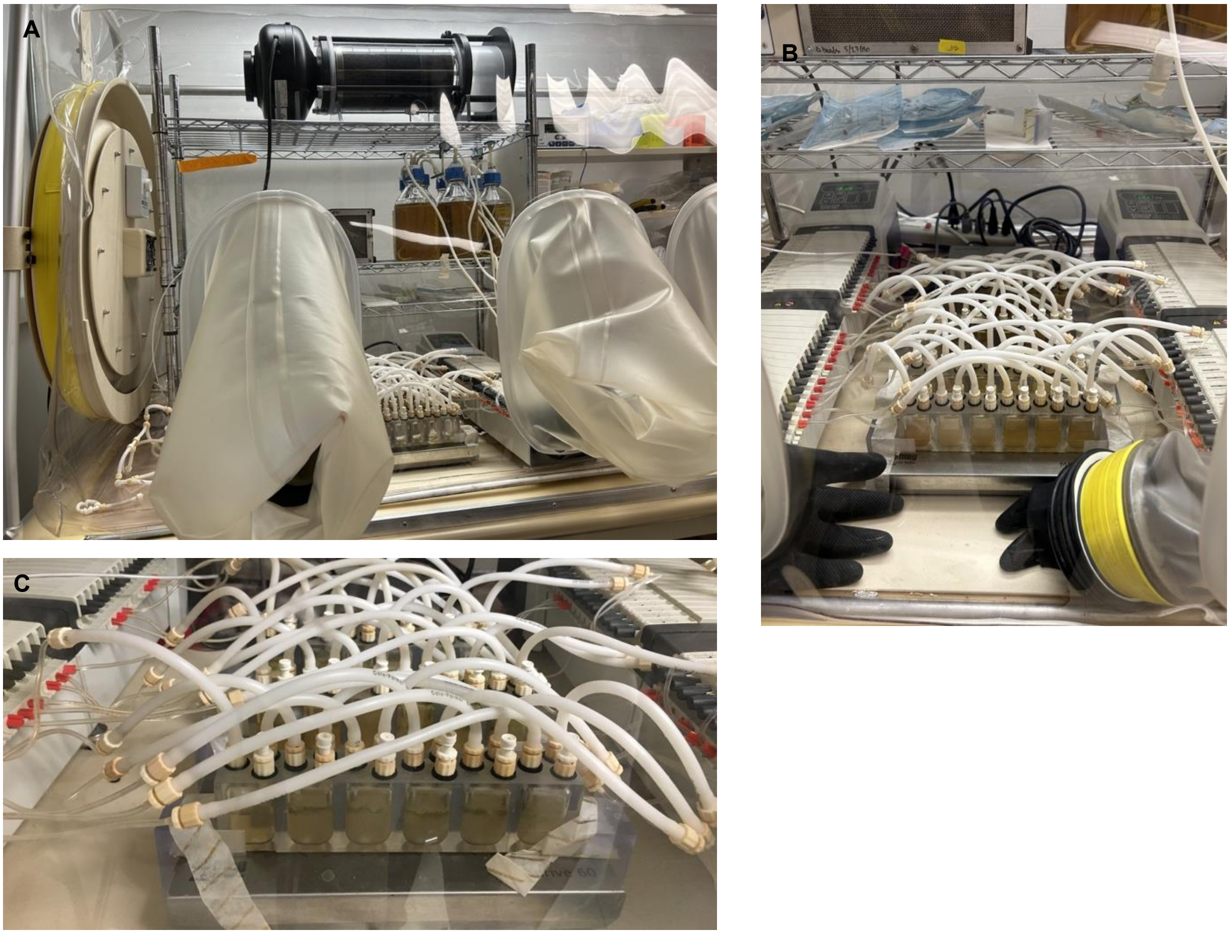


**Supplemental Figure 2. Photographs of miniature bioreactor arrays (MBRAs) set up in an anaerobic chamber**. (A) Anaerobic chamber with MBRAs, media bottles, and hydrogen sulfide removal column. (B) Closer view of pumps and MBRAs. (C) Close-up of MBRA strips on stir plate.

**Table S1. Characteristics of stool donors**

| Patient ID | Gender | Age | Race | Disease status |
|---|---|---|---|---|
| HC_1 | M | >50 | White | N/A |
| HC_2 | F | 61 | White | N/A |
| HC_3 | F | 40 | White | N/A |
| HC_4 | F | 43 | White | N/A |
| HC_5 | M | 30 | White | N/A |
| HC_6 | F | 38 | White | N/A |
| HC_7 | F | 43 | White | N/A |
| CD_1 | F | 33 | White | Quiescent |
| CD_2 | F | 59 | White | Active |
| UC_1 | F | 54 | White | Quiescent |
| UC_2 | M | 37 | Native Hawaiian/Pacific Islander | Quiescent |
| UC_3 | F | 40 | White | Quiescent |
| UC_4 | M | 55 | White | Quiescent |

M: Male, F: Female, N/A: Not applicable
